# *Rthoptera*: Standardised Insect Bioacoustics in R

**DOI:** 10.1101/2024.11.04.621915

**Authors:** Francisco Rivas, Cesare Brizio, Filippo M. Buzzetti, Bryan Pijanowski

## Abstract

1. Crickets were the first study subjects at the dawn of bioacoustics, yet, almost 100 years later, freely available and robust analysis tools and protocols are still needed.
2. We introduce *Rthoptera*, a new open-source R package for the analysis of insect acoustic signals, offering accurate graphics and standardised temporal and spectral measurements through a streamlined workflow. Our package delivers results in a fraction of the time typically required by multi-software methods. Most of the functions have interactive versions (Shiny applications), facilitating their adoption by researchers with any level of technical expertise.
3. New acoustic metrics are introduced: Spectral Excursion, Pattern Complexity Index, Temporal Excursion, Dynamic Excursion, and Broadband Activity Index.
4. Accompanied by an appropriate recording protocol, our tool can become the backbone of a standard analysis protocol for comparative insect bioacoustics.

## 1 INTRODUCTION

The current rate of species loss is outpacing our ability to track biodiversity change, highlighting the need for improved monitoring systems (O’Connor et al., 2020). Echoing the way in which satellite remote sensing revolutionised vegetation monitoring in remote areas (Huete, 2012), autonomous sound recorders are helping researchers track soniferous fauna at the community and ecosystem scales (Buxton et al., 2018). The study of animal sounds has followed two main frameworks. The soundscape ecology/ecoacoustics framework emphasizes large-scale patterns of biological acoustic activity over time and over ecological and disturbance gradients (Pijanowski et al., 2011; Pijanowski and Fuenzalida, 2024). Bioacoustic approaches usually focus on individual species to develop acoustic descriptions of species-specific signals, use them as keys to identify the species in new data, which are then manually annotated (Sugai et al., 2019) to finally be used in training automated detectors with the help of pattern matching algorithms (Hafner and Katz, 2018) and neural networks (Pijanowski et al., 2024). Although automatic identification of sounds has seen tremendous progress in birds (Kahl et al., 2021) and bats (Roemer et al., 2021), its development in insects is lagged. In the Global South, the required acoustic keys are not always available due to lack of bioacoustic training and/or insufficient resources to support the intensive field work required to record and collect voucher specimens before describing their acoustic signals (Riede, 2018). Due to these factors, basic insect bioacoustics is still a developing field in much of this region. This presents both challenges and opportunities for researchers in understudied areas, while also emphasising the need to cultivate a new community of practice. Such a community would accelerate the advancement of these promising tools, which have diverse applications ranging from agricultural pest monitoring (Mankin et al., 2000, 2011, 2021; Njoroge et al., 2017) to the use of sound-producing insects as proxies to monitor terrestrial and freshwater (Greenhalgh et al., 2021; Linke et al., 2018) ecosystems affected by climate change and habitat loss (Riede and Balakrishnan, 2024). Bioacousticians often use a multi-software approach to obtain measurements and generate graphs, resulting in time-consuming and tedious workflows. While free versions of bioacoustic software exist, they are typically optimised for analysing frequency-modulated signals, such as those produced by mammals and birds (Charif et al., 2010), and many tools are limited due to their proprietary nature. A popular alternative is open-source software like R (Ihaka and Gentleman, 1996; Decan et al., 2015), although it requires some programming skills. This caveat can be addressed by converting functions into a graphical user interface (GUI) using open source packages for web application development, such as Shiny (Jia et al., 2022). Here, we introduce a novel Shiny-powered package designed to facilitate both the extraction of taxonomically relevant spectral and temporal statistics and the production of standardised graphs commonly used in insect bioacoustics. Leveraging classic R packages for bioacoustics, *Rthoptera* allows researchers to interactively import, preprocess, and analyse insect sounds with fast extraction of both “classic” and novel acoustic metrics. Our workflow is semi-automated, minimising human bias, and returning results in a fraction of the time required by conventional methods using a multi-software approach. In addition to traditional static plots for printed publications, *Rthoptera* produces interactive plots in HTML format. The main goals of our work are 1) encouraging the production of high-quality analyses by researchers without requiring extensive expertise in R or bioacoustics and 2) providing reproducible, standardised, and interactive methods for producing graphs and measurements of insect acoustic signals. The latest version of *Rthoptera* can be freely downloaded and installed directly from *R* or *RStudio* interface using:

~~~
install.packages("remotes") remotes::install_github("naturewaves/Rthoptera")
~~~

## 2 PACKAGE DEVELOPMENT AND FEATURES

In this manuscript, the names of regular R functions are presented in monospaced font followed by parentheses (e.g., `function_name()`), Shiny apps are displayed in monospaced font without parentheses (e.g., `app_name`), and R packages are italicized (e.g., *packageName*). After consolidating the main processing, visualisation and data analysis techniques used by researchers, we built upon routines from *seewave* (Sueur et al., 2008) to create new plotting and analysis functions. Static plots are generated using *ggplot2* (Wickham, 2016), while interactive plots use *plotly* (Inc., 2015). After developing the functions, we used OpenAI ChatGPT-4 (OpenAI, 2024) as an assistant for coding the *Shiny* applications (Chang et al., 2024) and for developing the documentation and formal testing. Each code chunk generated in this way was manually analysed and debugged. Along with regular R functions for analysis and plotting, *Rthoptera* offers 16 Shiny interactive applications. Each of these apps can be invoked with a call to the `launchApp()` function. The package was tested manually and programmatically with *usethis* (Wickham et al., 2024) and *devtools* (Wickham et al., 2022), reaching 98% coverage, while operating system tests for Windows, MacOS, and Linux where successfully completed with *rhub* (Csárdi and Salmon, 2024). A summary of the functions and applications can be found in Table 1.

**Table 1.**
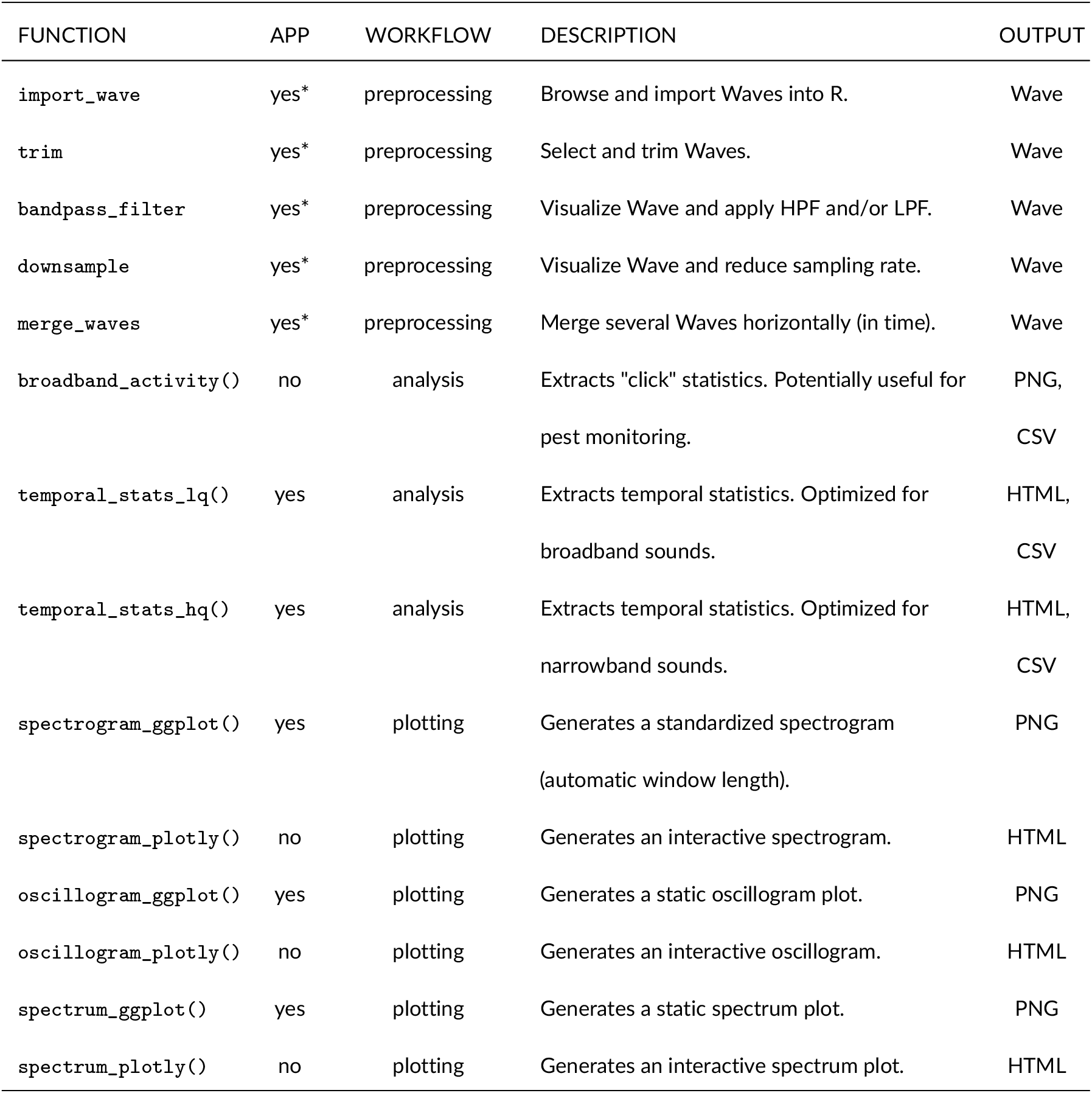
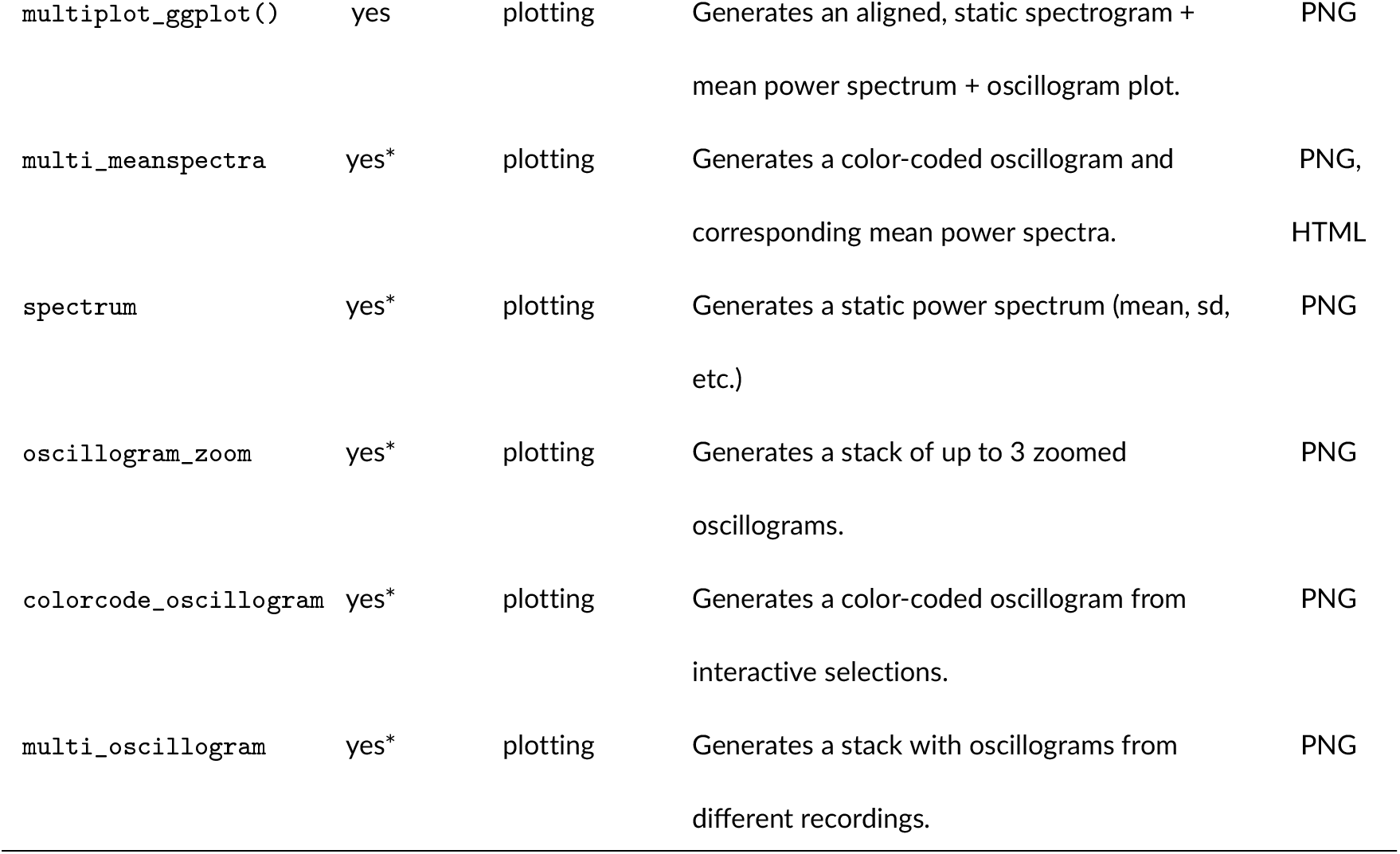
Summary of the functions in Rthoptera. *Available only as app. Apps should be called through the `launch_app()` function.

| FUNCTION | APP | WORKFLOW | DESCRIPTION | OUTPUT |
| --- | --- | --- | --- | --- |
| `import_wave` | yes* | preprocessing | Browse and import Waves into R. | Wave |
| `trim` | yes* | preprocessing | Select and trim Waves. | Wave |
| `bandpass_filter` | yes* | preprocessing | Visualize Wave and apply HPF and/or LPF. | Wave |
| `downsample` | yes* | preprocessing | Visualize Wave and reduce sampling rate. | Wave |
| `merge_waves` | yes* | preprocessing | Merge several Waves horizontally (in time). | Wave |
| `broadband_activity()` | no | analysis | Extracts "click" statistics. Potentially useful for pest monitoring. | PNG,<br>CSV |
| `temporal_stats_lq()` | yes | analysis | Extracts temporal statistics. Optimized for broadband sounds. | HTML,<br>CSV |
| `temporal_stats_hq()` | yes | analysis | Extracts temporal statistics. Optimized for narrowband sounds. | HTML,<br>CSV |
| `spectrogram_ggplot()` | yes | plotting | Generates a standardized spectrogram (automatic window length). | PNG |
| `spectrogram_plotly()` | no | plotting | Generates an interactive spectrogram. | HTML |
| `oscillogram_ggplot()` | yes | plotting | Generates a static oscillogram plot. | PNG |
| `oscillogram_plotly()` | no | plotting | Generates an interactive oscillogram. | HTML |
| `spectrum_ggplot()` | yes | plotting | Generates a static spectrum plot. | PNG |
| `spectrum_plotly()` | no | plotting | Generates an interactive spectrum plot. | HTML |
| `multiplot_ggplot()` | yes | plotting | Generates an aligned, static spectrogram + mean power spectrum + oscillogram plot. | PNG |
| `multi_meanspectra` | yes* | plotting | Generates a color-coded oscillogram and corresponding mean power spectra. | PNG,<br>HTML |
| `spectrum` | yes* | plotting | Generates a static power spectrum (mean, sd, etc.) | PNG |
| `oscillogram_zoom` | yes* | plotting | Generates a stack of up to 3 zoomed oscillograms. | PNG |
| `colorcode_oscillogram` | yes* | plotting | Generates a color-coded oscillogram from interactive selections. | PNG |
| `multi_oscillogram` | yes* | plotting | Generates a stack with oscillograms from different recordings. | PNG |

### 2.1 Demo Data

The companion package *RthopteraSounds* includes a small dataset of insect recordings made using diverse equipment and settings. These samples are used in the vignettes to illustrate the use of the apps and functions in *Rthoptera*. Additional recordings are used in the supplemental material to present a comparison with a “manual” methodology. These recordings were selected and plotted by authors 2 and 3 based on the manual measurements in Adobe Audition (Adobe Inc., 2024) and manual processing of screenshots with a generic image editor (Microsoft Corporation, 2024). The same files where parsed in *Rthoptera* to obtain graphs and measurements which were then compared to the traditional approach. Example recordings in this dataset are available in a companion package which can be installed using:

~~~
remotes::install_github("naturewaves/RthopteraSounds")
~~~

### 2.2 Analysis formats

Insect acoustic signals are usually measured and visualised in three basic analysis formats:

- **Oscillogram**: A waveform representing changes in the amplitude (energy) of a sound over time, fluctuating around zero. Researchers often use an oscillogram of the entire calling song along with a zoomed-in portion of one of the smaller structures to illustrate the temporal patterns. All temporal measurements in *Rthoptera* are calculated on the envelope of the waveform, which is extracted with the `env()` function from *seewave*.
- **Mean Power Spectrum**: A vector representing the distribution of power along the frequency axis, often illustrated as a frequency versus amplitude plot, from which all spectral measurements in *Rthoptera* are derived. While other software allows visualising the mean spectrum of single time frames (i.e., “instantaneous spectrum”), our package avoids this approach by design. Instead, we consider the complete `Wave` object, providing a more representative profile of the signal that can be reliably referenced to the corresponding spectrogram and oscillogram plots.
- **Spectrogram**: A colour-coded (or grey-scale) plot derived from a matrix representing the intensity of sounds across frequency (rows) and time (columns), which is the result of a Short-Time Fourier Transform (STFT) applied to the `Wave` object. Also referred to as a “sonogram”. This plot is often used for visualisation purposes and to calculate some spectral acoustic indices (Sueur et al., 2014). Extracting “manual”, “point” measurements (e.g., peak frequency, start and end of sounds, etc.) directly from the spectrogram is not recommended due to the loss of time resolution intrinsic in the STFT.

### 2.3 Terminology for temporal units

Nomenclature inconsistencies in the literature motivated Baker and Chesmore (2020) to develop a standardisation of the bioacoustic terminology for insects. Some of these inconsistencies are still difficult to resolve, mainly because there is no clear categorisation to distinguish between terms linked to behaviours (e.g., hemisyllable, syllable) and those purely acoustic (e.g., pulse train). Considering that researchers may utilise our package for diverse purposes, we opted for using a neutral terminology, leaving the “translation” of the appropriate terms up to their best judgement:

- **Peak**. The `temporal_stats_lq()` function identifies individual peaks in the envelope, usually corresponding to “tooth impacts”. See Baker and Chesmore (2020) for details.
- **Train**. A set of consecutive peaks, which can be translated as “pulse” or “syllable”, depending on the taxon or sound unit under study. For authoritative definitions of those terms and the underlying physiology, see Bennet-Clark and Bailey (2002); Gerhardt and Huber (2002); Walker and Carlysle (1975); Huber et al. (1989); Otte (1992). “Train” is used in both temporal statistics functions.
- **Motif**. Any of the aggregations of trains, including “dyplosyllable”, “echeme”, “trill”, etc.). This term is used in both temporal statistics functions. The correct translation will depend on the taxon and grouping parameter values.

## 3 PREPROCESSING

Although high signal-to-noise ratio (SNR) recordings can be achieved under controlled conditions, noise is a common issue in field recordings, requiring preprocessing of the sound files before analysis. *Rthoptera* offers a set of interactive functions (Shiny apps) to perform the most common audio preprocessing tasks in bioacoustics.

### 3.1 Import

While importing audio files into memory traditionally involves typing at least two commands into the console or using the file menu including `readWave` (Ligges et al., 2023), the `import_wave` app allows the user to browse the computer drives and import audio with the added convenience of listing only sound files (WAVE, WAC or MP3).

### 3.2 Downsample

When a signal of interest (SOI) occurring within the audible range (e.g., < 18 kHz) is recorded with ultrasonic equipment at a very high sampling rate (e.g. 250 kHz), much of the higher portion of the spectrum is irrelevant for analysis or plotting. In such cases, and unless the researcher is interested in very faint ultrasonic frequencies (Brizio, 2023), it is convenient to downsample the audio file to improve computational performance. The `downsample()` allows one to inspect an interactive mean power spectrum to evaluate the distance between the maximum frequency of interest (MaxFOI) and the Nyquist frequency (i.e., the maximum recorded frequency, equivalent to half the sampling frequency) before deciding to downsample. Depending on the recording bandwidth, available downsampling values (“new Nyquist”) are constrained to 48, 96 and 125 kHz.

### 3.3 Band-pass filter

The `band_pass_filter()` allows the user to visually inspect an interactive spectrogram and mean power spectrum of a selected `Wave` object to assess the frequency range of the signal of interest (SOI) and identify the presence and range of non-target sounds (e.g., low-frequency noise and/or non-target higher-frequency signals) before applying a proper combination of high-pass-(HPF) and low-pass filter (LPF), as needed.

### 3.4 Trim

The `trim()` facilitates storing selections of a `Wave` in the R environment by assigning a new name to create a separate object or reassigning the original name to overwrite it. This app can also be used to extract individual sound types (e.g., ‘opening strokes’) that can be later merged into a single wave for separate analysis.

### 3.5 Merge

The `merge_waves()` allows users to select several `Wave` objects from the R environment and merge them into a new object, which can also be exported as a WAVE file. This is useful when one wants to obtain a representative measure of one element type such as the “opening stroke” (or opening hemisyllable), or the “closing stroke”.

## 4 ANALYSIS

### 4.1 Spectral Statistics

The `spectral_stats()` (Figure 2) performs automatic calculations of several acoustic features in the frequency spectrum. The function returns an interactive plot and a table with the results and the parameters used for the analysis. Although the plot is intended mainly to monitor the measurements and assist in parameter selection, we encourage researchers to download the HTML file and make it available as supplemental material for the readers to explore. The measurements returned in the table include:

**Figure 1.**
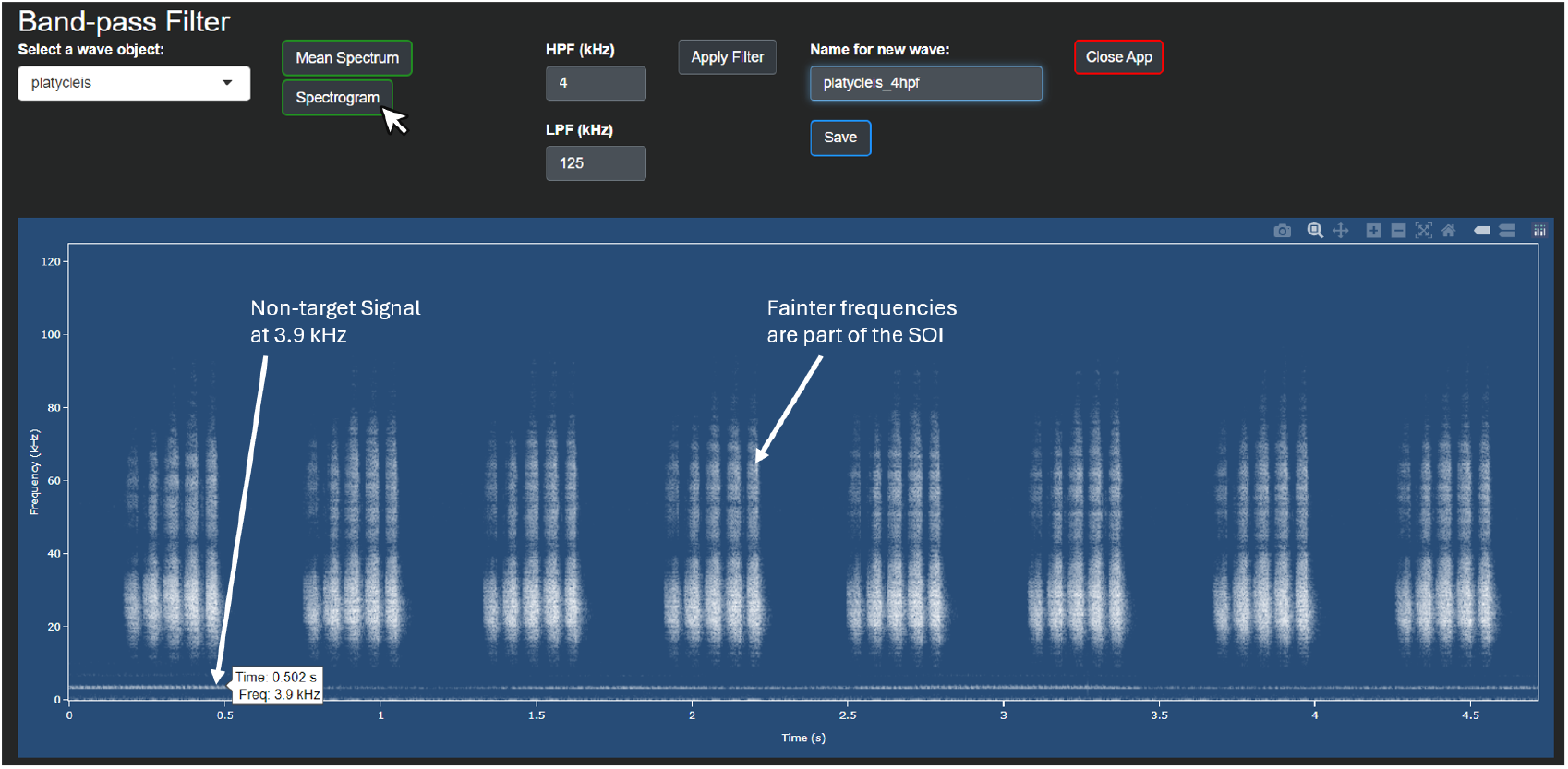
The `band_pass_filter()`, spectrogram view. The continuous signal around 3.9 kHz is the song of a different species. Since this is not part of the SOI, a 4 kHz HPF is needed. We can also see that the fainter frequencies between ≈ 40 and ≈ 80 kHz are part of the SOI, so no LPF is required.

**Figure 2.**
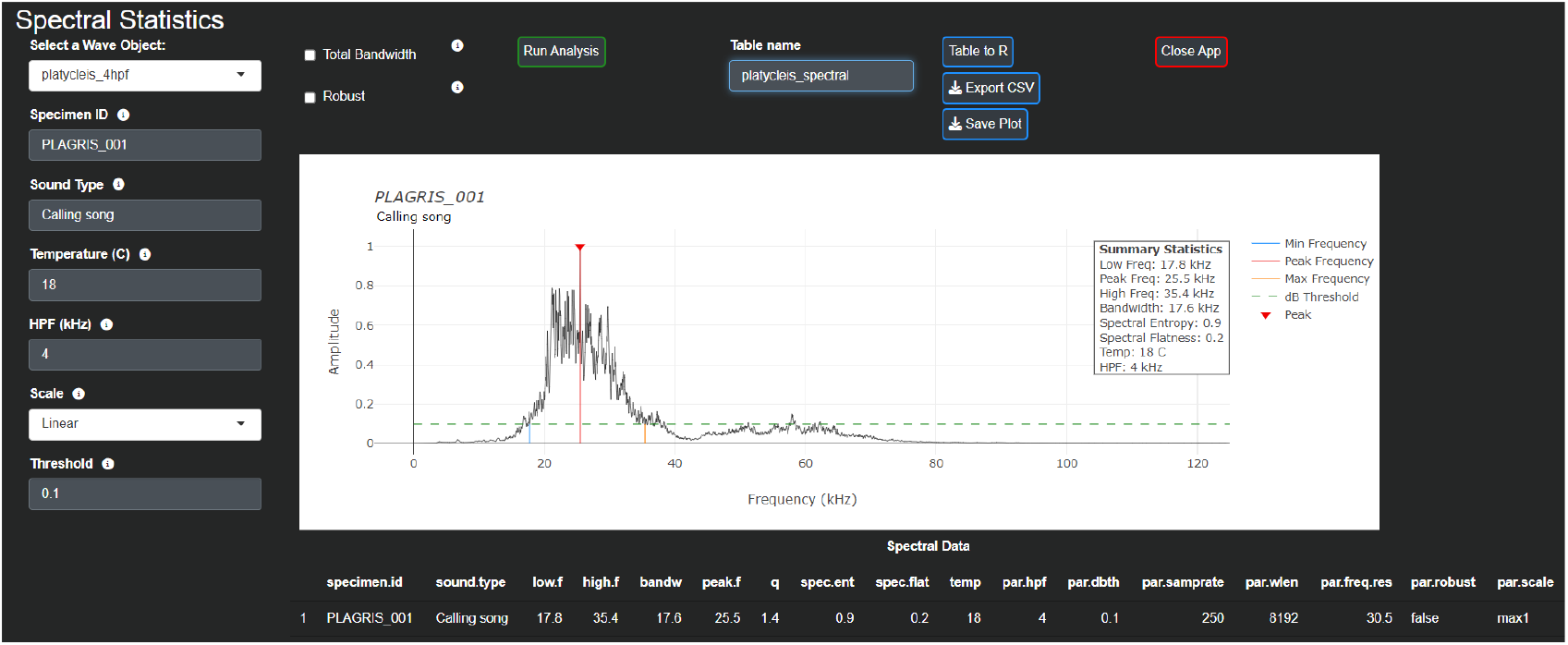
The `spectral_stats()` returns quick, automatic spectral measurements.

- **Peak Frequency**: The frequency with maximum amplitude value. Also known as “carrier frequency”.
- **Low Frequency**: The first frequency below the selected amplitude threshold to the left of the peak frequency (assuming that the frequency is on the x-axis).
- **High Frequency**: The first frequency below the selected amplitude threshold to the right of the peak frequency (assuming that the frequency is on the x-axis).
- **Spectral Excursion**: The length of the spectral contour along the frequency axis between the Low Frequency and the High Frequency.
- **Spectral Entropy**: The Shannon entropy of the spectrum, calculated with `sh()` function from *seewave*.
- **Spectral Flatness**: The spectral flatness of the `Wave`, calculated with the `sfm()` function from *seewave*.
- **Quality Factor**: The Quality Factor (Q) is defined as the peak frequency divided by the bandwidth and it was incorporated experimentally in *Rthoptera*. In rigour, the Q factor is equivalent to that described in the classic literature only when the threshold is set at −3dB or −10 dB. See Bennet-Clark (1999) for advice on its use and interpretation.

For publication-grade static plotting of the power spectrum, *Rthoptera* provides two dedicated applications: `spectrum_plot()` for a single spectrum and `multi_meanspectra()` for overlayed mean spectra from multiple selections.

### 4.2 Temporal Statistics

Accounting for the diversity of sound production in insects, *Rthoptera* offers two functions to extract temporal statistics from insect sound recordings. For practical reasons, we distinguish between two call types in insects, High-Quality Factor (HQ) and Low-Quality Factor (LQ) following Montealegre-Z and Morris (1999). HQ calls have a tonal, “musical” quality, producing a narrow band usually accompanied by one or several harmonics of varying intensity. HQ calls are produced by most field crickets, ground crickets, and tree crickets. For these signals, it is appropriate to use a cutoff amplitude threshold to find the start and end of each sound, which is the approach used by the `temporal_stats_hq()` (Figure 3) function. In contrast, LQ calls have a wide-band frequency profile, with a harsh, “noisy” timbre, common in most katydids, bush crickets, grasshoppers, and coleopterans. The `temporal_stats_lq()` function (Figure 4) uses a ‘peakfinder’ procedure to detect short local maxima along the waveform, allowing for the accurate measurement of transient (LQ) sounds and trains that are too faint compared to the loudest parts of the call and therefore are harder to capture with the amplitude threshold approach. See Table 2 for guidance on the use of temporal statistics functions for each of the main soniferous taxonomic groups.

**Table 2.**
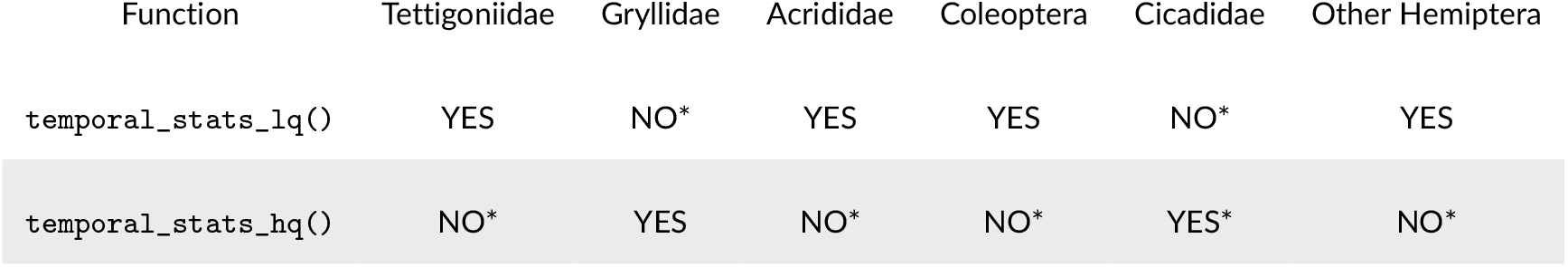
Suitability of each temporal statistics function for six taxonomic groups known to produce sounds. * Although the function is not optimised for this taxon, it can still be used if appropriate parameters are chosen. Check the vignettes for guidance on this topic.

**Figure 3.**
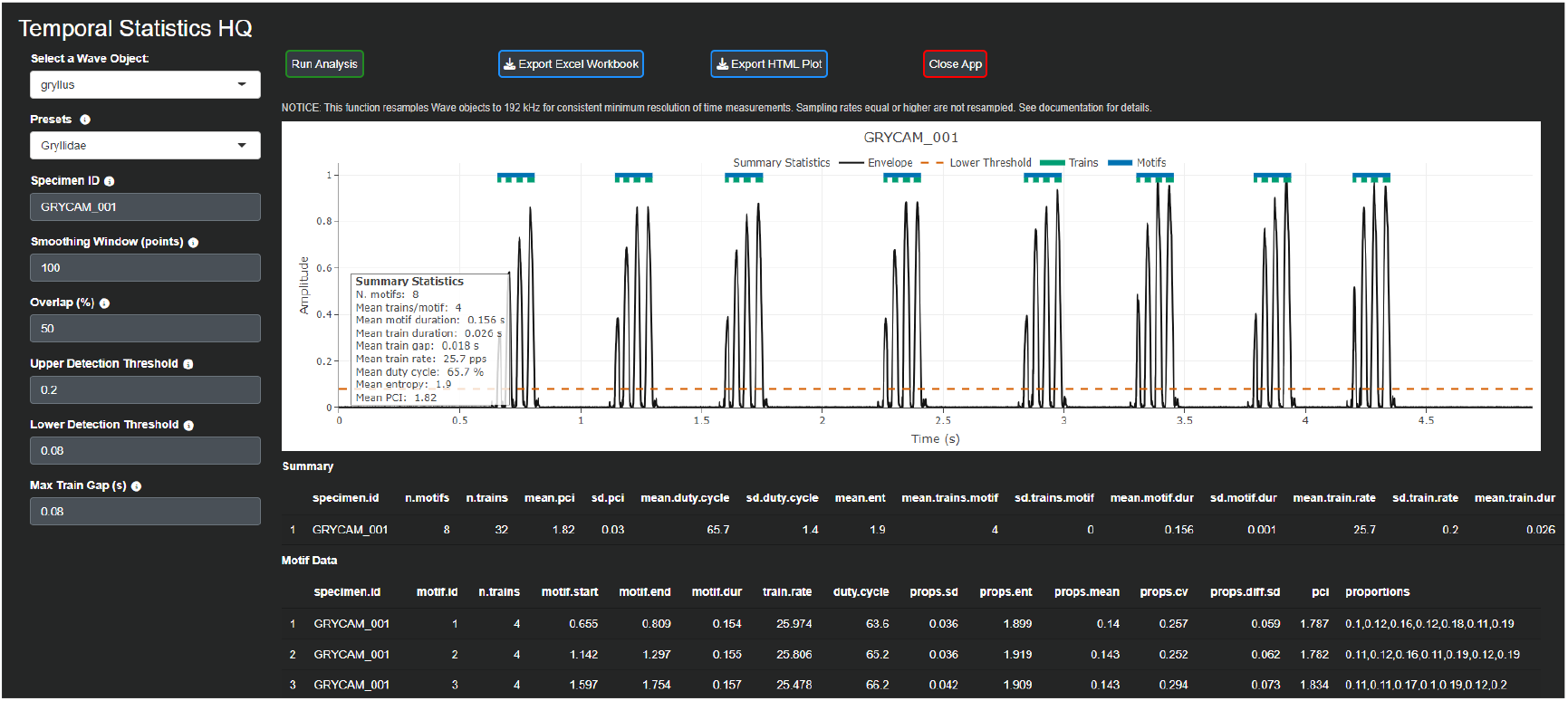
texttttemporal_stats_hq() calculates automatic temporal measurements, including the novel Pattern Complexity Index. This function is optimised for tonal calls, like this example of *Gryllus campestris*.

**Figure 4.**
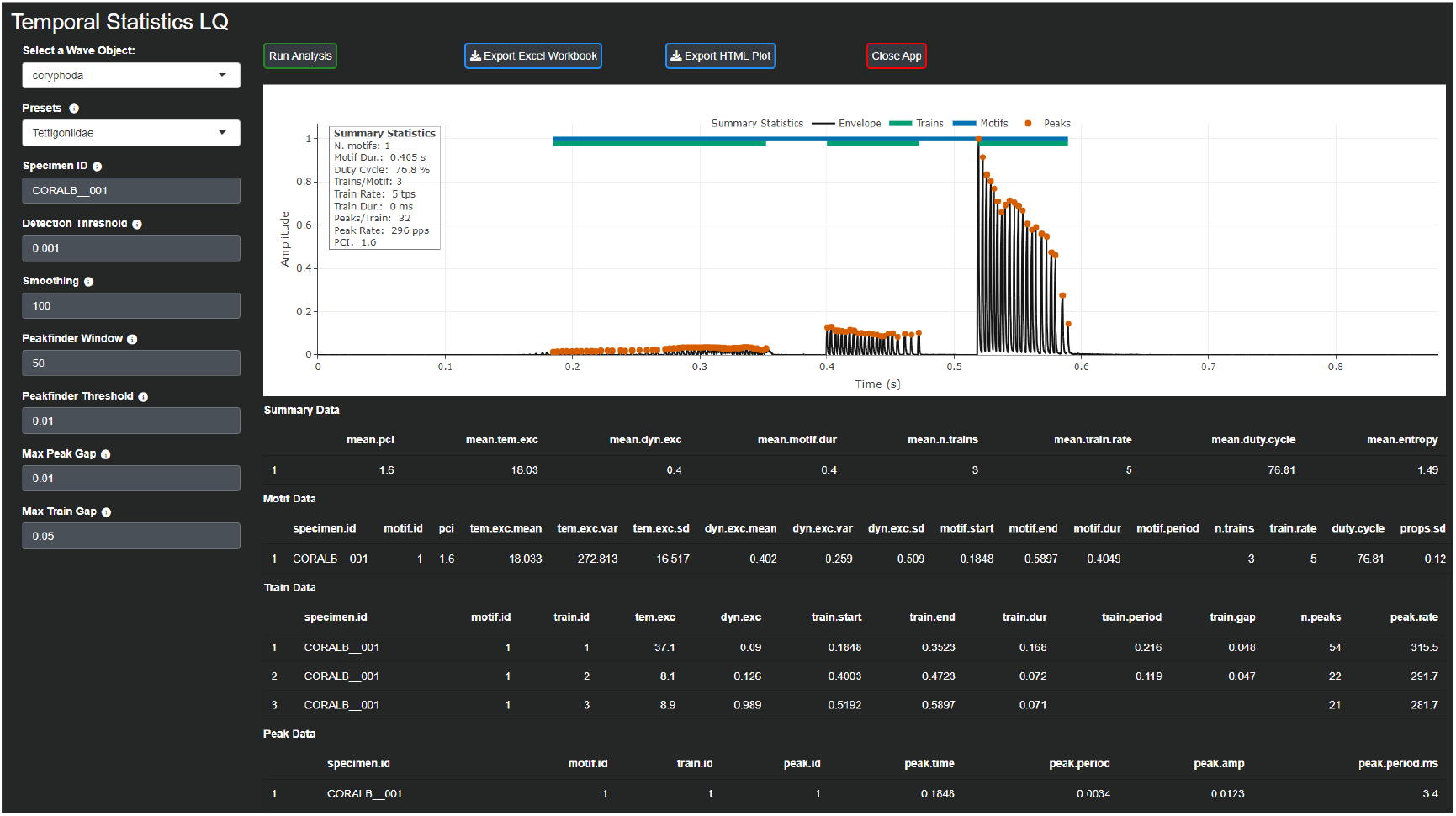
`temporal_stats_lq()` detects local peaks (tooth impacts), accurately measuring “transient”. This example showcases a single song of *Coryphoda albidicollis*.

*Rthoptera* introduces three novel temporal metrics: the Pattern Complexity Index (PCI), Temporal Excursion (TE), and Dynamic Excursion (DE). The PCI aims to characterise the temporal structure of a motif (e.g., an echeme, a calling song, etc.) with a metric that is robust to the natural variability caused by temperature differences during recording. To achieve this, we take the proportions of the duration of trains and gaps across the motif and then calculate PCI as follows:

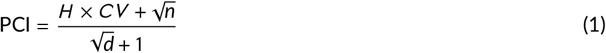

where *H* is the Shannon entropy of the proportions, *CV* is the coefficient of variation of the proportions, *n* is the number of trains in the motif and *d* is the duration of the motif. The resulting PCI is rounded to three decimal places. TE measures the variability in ‘acceleration’ within each train by taking the absolute differences between consecutive peak periods in the train and adding them together. Consequently, a train containing regularly-spaced peaks will return a low TE value. TE is calculated as follows:

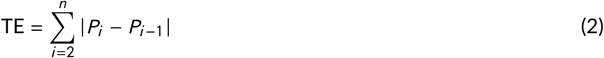

where *P*_*i*_ represents the *i*-th peak period, in milliseconds, and *n* is the total number of peaks for a given train. Missing values are ignored in the summation. Along with PCI, measures of dispersion in TE across trains could help characterise overall temporal complexity of the motif. Similarly, the Dynamic Excursion captures the variability in amplitude of peaks within a train by taking the absolute differences between consecutive peak amplitudes in a train and adding them together. In species emitting sound in both wing strokes, the closing hemisyllable generally yields a higher DE value than the opening one. DE is calculated as follows:

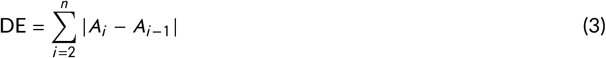

where *A*_*i*_ represents the amplitude of the *i*-th peak relative to the maximum amplitude in the analysed `Wave`, and *n* is the total number of peaks for a given train. Missing values are ignored in the summation. Figure 5, shows that the first train (hemisyllable) is the faintest in the motif, with the lower DE value and the highest TE value, conveying potentially relevant information about the identity of the signaller. Furthermore, TE and DE values could be useful for automatic train classification in some species, expediting batch analysis in future deployments of the package and complementing taxonomic studies.

**Figure 5.**
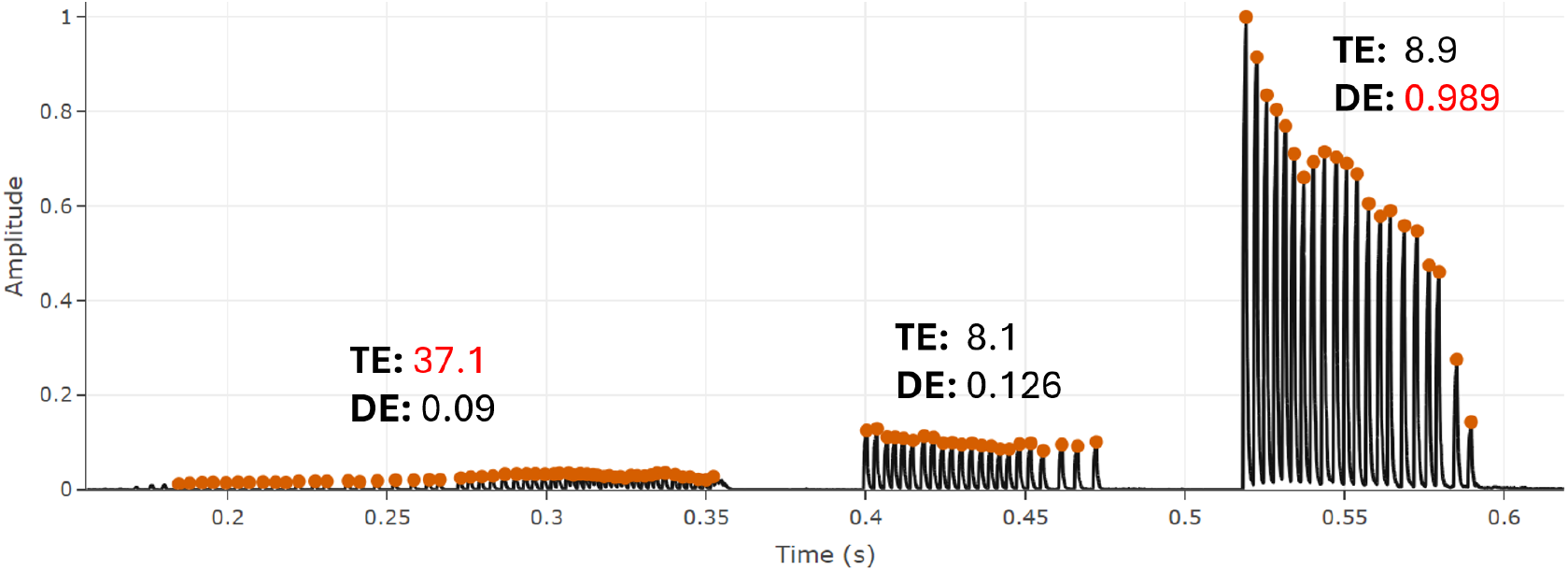
Temporal and Dynamic Excursion indices values of a *Coryphoda albidicollis* obtained with the `temporal_stats_lq()`. Extreme values for each index are highlighted in red.

Both temporal statistics applications return a set of tables, including summary, motif data, train data, and the parameters used for the analysis, which can be downloaded as a single Excel Workbook. TE and DE are reported in the “Train_Data” table from the `temporal_stats_lq()` function and app.

The summary table consists of the following metrics:

- **Number of motifs**: The total number of motifs (e.g., ‘echemes’) detected in the recording.
- **Number of trains**: The total number of trains (e.g., ‘hemisyllables’) detected in the recording. The following metrics are reported as mean and standard deviation values calculated across motifs in the `Wave` object:
- **PCI**: The Pattern Complexity Index value.
- **Duty Cycle**: A measure of the ratio between the above-threshold duration and the total duration of the motif.
- **Entropy**: The Shannon Entropy calculated over the proportions of trains and gaps within the motif. Only the mean is reported.
- **Trains per Motif**: The number of trains in each motif.
- **Motif duration**: The duration of the motifs.
- **Train rate**: The mean number of trains per second within a motif.
- **Train duration**: The mean duration of trains in seconds.
- **Train period**: The mean period of trains. This is calculated as the distance from the start of the train to the start of the next train in seconds.
- **Gap duration**: The mean duration of gaps in seconds.

## 5 PLOTTING

*Rthoptera* offers a suite of apps designed for standardised plotting, tailored to meet the needs of researchers in insect bioacoustics. These apps provide tools for generating the most commonly utilised graph types in the field, including oscillograms, power spectra, and spectrograms. The standardised approach of the functions facilitates consistency and reproducibility across studies. All colour-coding functions use colourblind-safe palettes, ensuring accessibility and clarity in visualizations for all users.

- **Power Spectrum**. Two apps are available for this plot type:
  - `spectrum_plot()`. This app generates a customizable Power Spectrum plot (Figure 7) with ‘mean’, ‘median’, ‘var’ (variance), and ‘sd’ (standard deviation) functions available to summarize the spectrum. The user can also change the orientation of the axes and limit the frequency range.
  - `multi_meanspectra()`. This app allows to select different sections from an oscillogram and add them to a Mean Power Spectra Plot, where each selection is automatically colour-coded and added upon the previous one. This allows to visually assess the spectral profile of smaller units in a calling song. The `multi_meanspectra_ggplot` app produces a static version (Figure 8) of the mean power spectra, which is better suited for printed publications.
- **Oscillogram**. This essential plot in insect bioacoustics can be generated in three different forms in the package:
  - `oscillogram_zoom`. This app allows the user to stack two zoomed waveforms below the main (e.g., a calling song) oscillogram.
  - `oscillogram_interactive()`. This function creates an interactive oscillogram which can be saved in HTML format, which can be made available in online repositories for free user exploration.
  - `multioscillo`. This app allows to stack oscillograms of different `Wave` objects, facilitating visual comparison between species or different calls of a single species. The user can choose between showing the x-axis or replace it with a scale bar.
  - `colorcode_oscillogram`. This app allows to colour-code selections of a spectrogram. An example use for this functionality is to highlight different individuals (e.g., male and female duet) in a recording.
- **Spectrogram**. Spectrograms (or ‘sonograms’) are frequently used in bioacoustics of most taxa. In insects, however, researchers often struggle to find the optimal parameters which would yield a clear image with an acceptable trade-off between spectral and temporal resolutions. The `spectrogram()` solves this by automatically selecting a window length (WL) accounting for the sampling rate and duration of the `Wave` as follows:

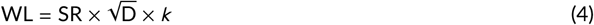

where SR is the sample rate, D is the duration of the `Wave` object, and *k* is a constant (20 × 10^−4^). Additionally, the app includes a “noise floor” parameter, which clips the cells below this threshold (in dB Full Scale) to remove background noise. Optionally, a flipped Mean Power Spectrum can be added on the right side of the spectrogram, sharing the Y-axis. The Power Spectrum can be plotted in dB or linear scale, both with reference to the local peak amplitude.
- **Multiple plot**. It is often convenient to plot all the features of a `Wave` simultaneously. The `multiplot()` (Figure 6) generates a standard spectrogram with a mean power spectrum (the same spectrogram described above) and adds an oscillogram on the bottom (sharing the x-axis), all in one click if the default parameters are used.

**Figure 6.**
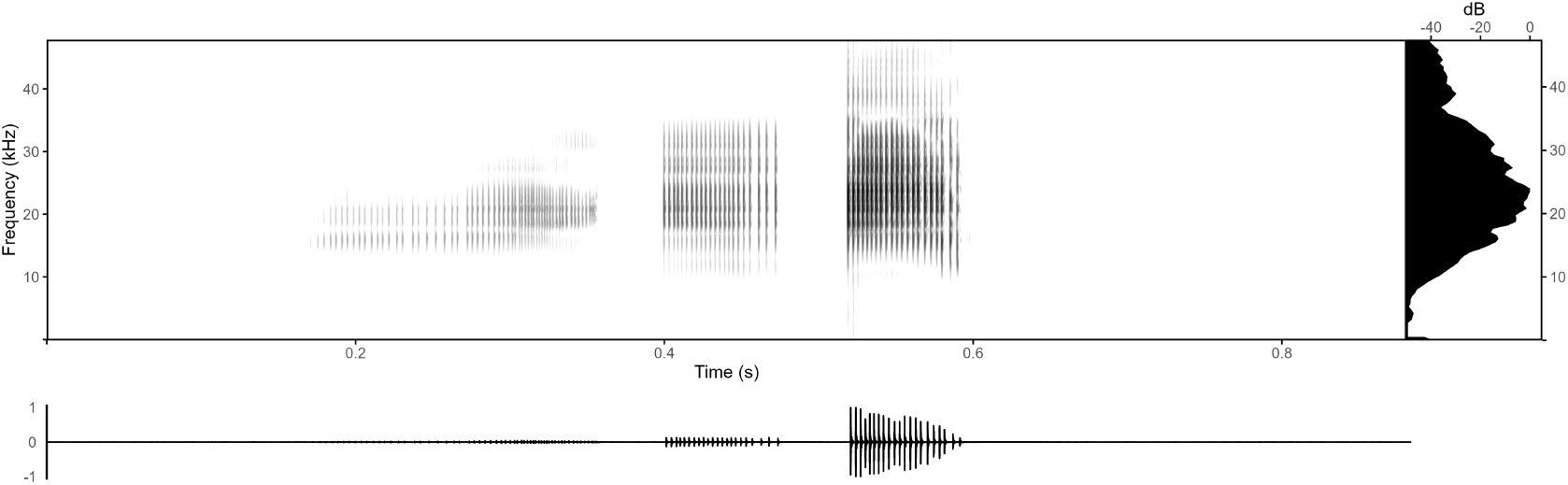
Standard “multiplot” of a single calling song of *Coryphoda albidicollis* obtained with one click using the default parameters in textttmultiplot().

**Figure 7.**
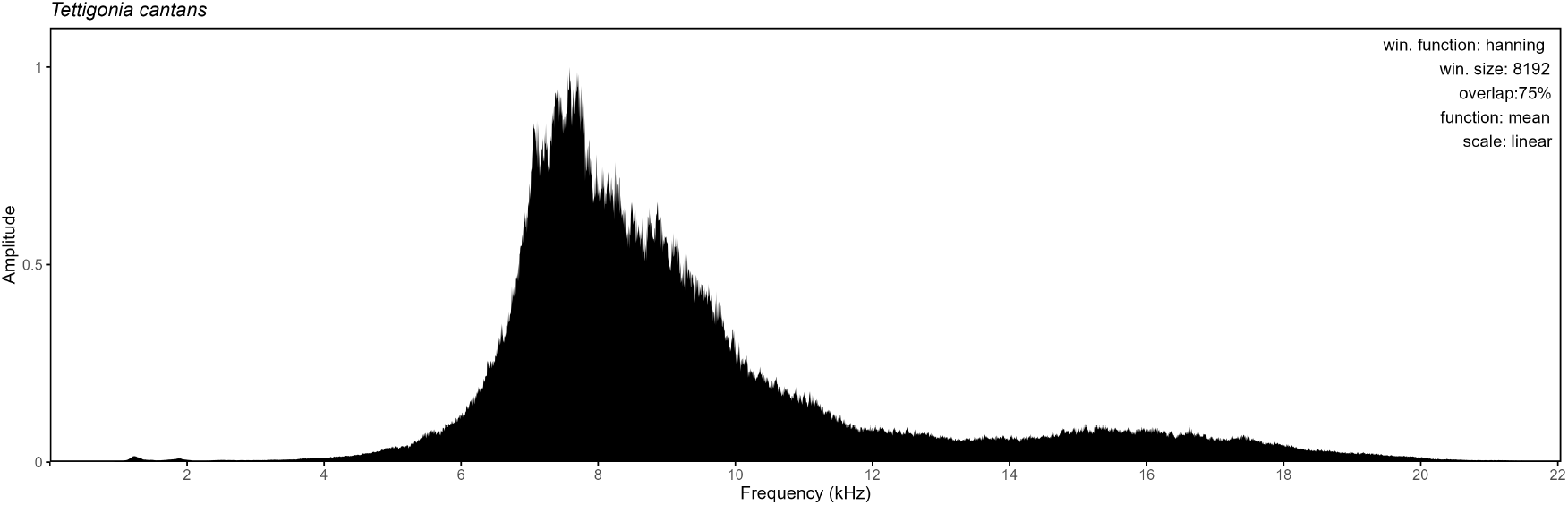
Mean Power Spectrum of a single calling song of *Tettigonia cantans* obtained with the `spectrum_plot()`. Including the parameters in the plot is optional and can be toggled off from a checkbox in the app.

**Figure 8.**
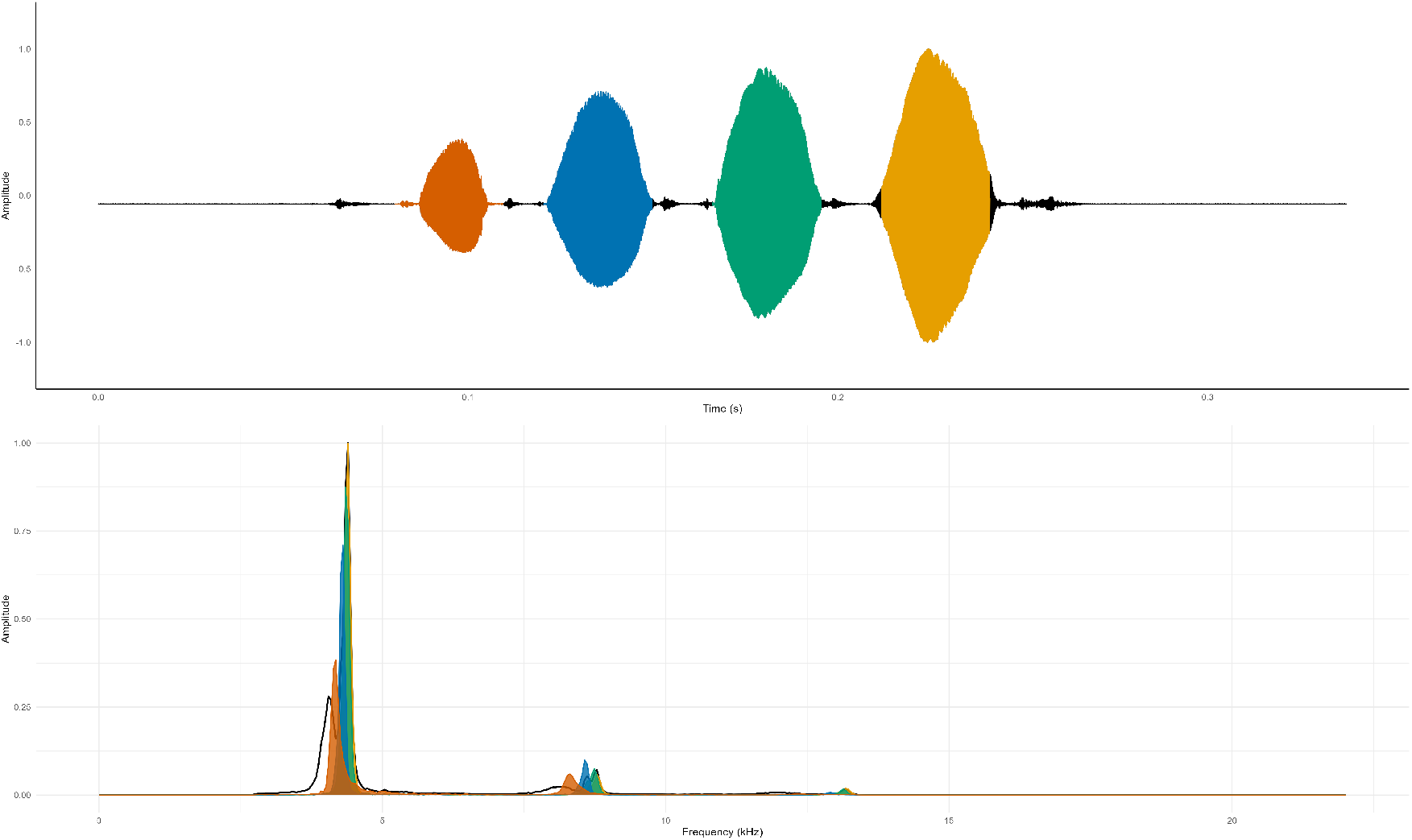
Output of the `multi_meanspectra_static()`. Individual pulse selections from *Gryllus campestris* are depicted.

## 6 POTENTIAL FOR PEST MONITORING

The `broadband_activity()` function creates a noise-reduced spectrogram and detects “clicks” (i.e, broadband, transient noises), extracting summary statistics including number of clicks, click height, and click centroid (kHz). The Broadband Activity Index (BBAI) corresponds to the proportion of elements in the spectrogram matrix (or cells in the spectrogram) which are classified as part of a click. Originally developed for analysing soundscape recordings to detect noise sources, it has potential for monitoring non-stridulating insect pests in stored food and wood, which incidental sound patterns and challenging recording conditions result in audio files that are not suitable for the other analytical approaches presented in this manuscript. Figure 9 shows an example of click detection with BBAI for a recording of Maize Weevil *Sitophilus zeamais* adults feeding on maize.

**Figure 9.**
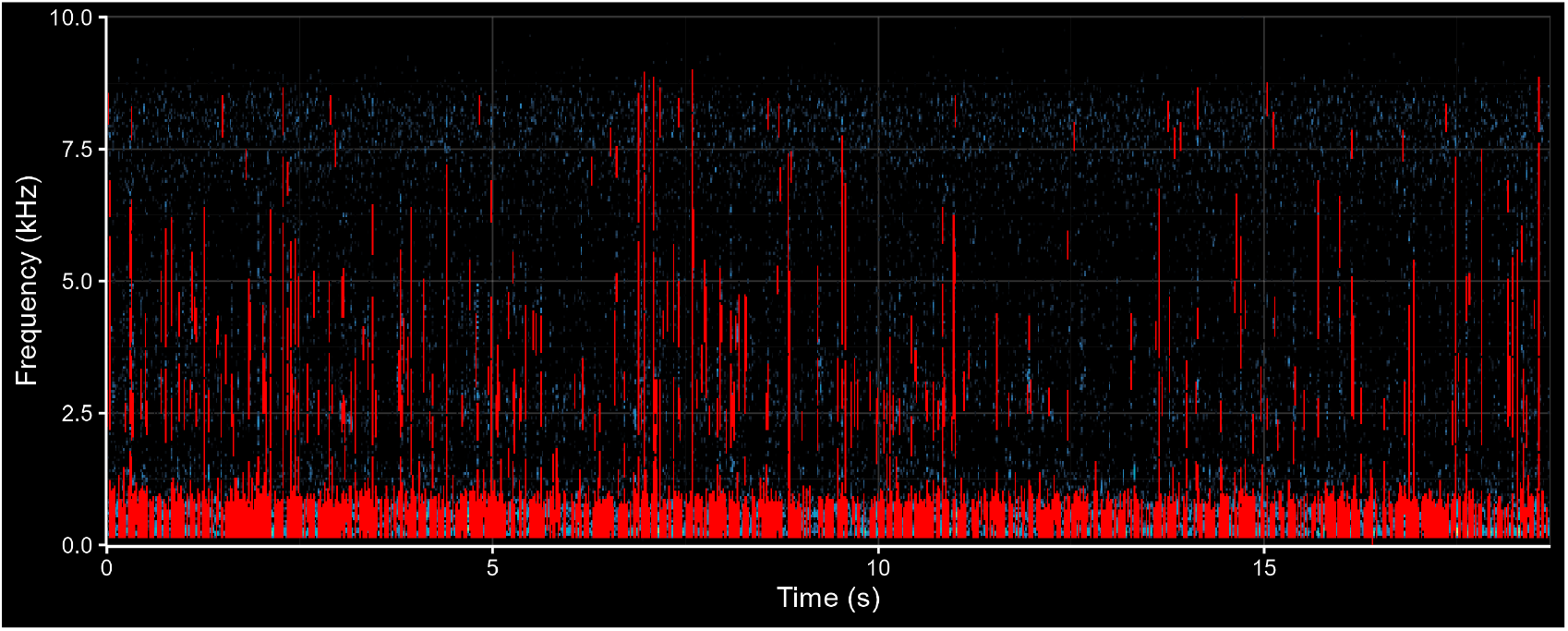
Plot obtained with the `broadband_activity()` function from a recording of *Sitophilus zeamais* feeding on maize. The “clicks” produced by the feeding activity are highlighted in red, while background noise is in light blue.

## 7 DISCUSSION

Insects are important indicators of ecosystem health and agricultural safety, with many species producing sound in terrestrial and aquatic (Aiken, 1985) habitats. Although acoustic monitoring has improved greatly in the last decades, automatic detection for insect species is lagged due to lack of baseline data. As a contribution to this effort, the *Rthoptera*, package offers an expedite and standardised workflow for insect bioacoustic analysis, facilitating the description of new bioacoustic types as well as the redescription of older ones by leveraging new techniques. These descriptions are the first step in a broader acoustics-based conservation endeavour, contributing to monitoring and conservation of habitats, communities, and species, and pest control. Our new approach for signal delimitation in broadband stridulations and the new metrics for temporal and dynamic variability can enrich the bioacoustics toolbox by considering dimensions that are otherwise overlooked. We hope that our package encourages researchers with any degree of expertise to conduct novel bioacoustics research. Our team is open to contributions to expand the capabilities of the package by suggesting new functions and fixes through the Issues section of the GitHub website.

## AUTHOR CONTRIBUTIONS

Author 1 conceived the package, developed the code, and wrote the manuscript. Author 2 co-designed the functionalities of the shiny applications, performed rigorous tests against various bioacoustic samples of his authorship, and contributed significantly to the manuscript. Author 3 provided expert feedback at each stage of development. Author 4 provided consistent guidance and funding throughout the process. All co-authors provided detailed reviews of the manuscript and approved its final version.

## ACKNOWLEDGEMENTS

We thank Klaus Riede, Baudewijn Odé, and Laurel Symes for their useful feedback in the early stages of development and Richard Mankin for facilitating his recording of *Sitophilus zeamais*. Author 1 was supported by National Aeronautics and Space Administration (NASA, USA) Award: 80NSSC21K1146, and the National Agency for Research and Development (ANID, Chile) Scholarship ID: 72210036. We dedicate this paper to the memory of Gianni Pavan, whose legacy inspired our work.

## DATA ACCESSIBILITY

The source code, vignettes and data used for demonstrations can be found on GitHub: https://github.com/naturewaves/ `Rthoptera` and the package’s website https://naturewaves.github.io/Rthoptera/.

